# A Monte Carlo method to estimate cell population heterogeneity

**DOI:** 10.1101/758284

**Authors:** Ben Lambert, David J. Gavaghan, Simon Tavener

## Abstract

Variation is characteristic of all living systems. Laboratory techniques such as flow cytometry can probe individual cells, and, after decades of experimentation, it is clear that even members of genetically identical cell populations can exhibit differences. To understand whether variation is biologically meaningful, it is essential to discern its source. Mathematical models of biological systems are tools that can be used to investigate causes of cell-to-cell variation. From mathematical analysis and simulation of these models, biological hypotheses can be posed and investigated, then parameter inference can determine which of these is compatible with experimental data. Data from laboratory experiments often consist of “snapshots” representing distributions of cellular properties at different points in time, rather than individual cell trajectories. These data are not straightforward to fit using hierarchical Bayesian methods, which require the number of cell population clusters to be chosen *a priori*. Here, we introduce a computational sampling method named “Contour Monte Carlo” for estimating mathematical model parameters from snapshot distributions, which is straightforward to implement and does not require cells be assigned to predefined categories. Our method is appropriate for systems where observed variation is mostly due to variability in cellular processes rather than experimental measurement error, which may be the case for many systems due to continued improvements in resolution of laboratory techniques. In this paper, we apply our method to quantify cellular variation for three biological systems of interest and provide Julia code enabling others to use this method.

## 2 Introduction

Variation, as opposed to homogeneity, is the rule rather than exception in biology. Indeed, without variation, biology as a discipline would not exist, since as evolutionary biologist JBS Haldane wrote, variation is the “raw material” of evolution. The Red Queen Hypothesis asserts organisms must continually evolve in order to survive when pitted against other - also evolving - organisms [1]. A corollary of this hypothesis is that multicellular organisms should evolve cellular phenotypic heterogeneity to allow faster adaptation to changing environments, which may explain the observed variation in a range of biological systems [2]. Whilst cell population variation can confer evolutionary advantages, it can be costly in other circumstances. In biotechnological processes, heterogeneity in cellular function can reduce yields of biochemical products [3]. In human biology, variation across cells can enable pathologies to develop; it can also frustrate treatment of illness because key subpopulations are missed by medical interventions that target “average” cell properties. For example, cellular heterogeneity helps some cancerous tumours to persist [4] and can make tumours more likely to evolve resistance to chemotherapies [5]. To discern whether observed variation is benign or requires remedy, methods of analysis are needed that can quantify and help to understand its source.

Mathematical models are essential tools for understanding cellular systems, whose emergent properties are the result of a nexus of interactions between actors. Perhaps the simplest flavour of mathematical model used in biological systems is an ordinary differential equation (ODE) that aggregates individual actors into compartments according to structure or function, and seeks to model the mean behaviour of each compartment. Data from population-averaged experimental assays can determine whether such models faithfully reproduce system behaviours and can be used to understand the structure of complex metabolic, signalling and transcriptional networks. The worth of such “population average” ODE models depends on whether averages mask substantial differences in individual behaviour [6]. In some cases, differences in cellular protein abundances due to biochemical “noise” are not biologically meaningful [7] and the system is well described by average cell behaviour. In others, there are functional consequences. For example, a laboratory study demonstrated that subpopulations of clonally-derived hematopoietic progenitor cells with low expression of a stem cell marker, diverged into a separate blood lineage from those with high expression [8].

Many modelling frameworks are available to describe cell population heterogeneity, with each posing different challenges for parameter inference. A recent review is presented in [9]. These approaches include modelling biochemical processes stochastically, where properties of ensembles of cells are represented by probability distributions that evolve according to chemical master equations. See [10] for a tutorial on stochastic simulation of reaction diffusion processes. Alternatively, population balance equations (PBEs) are typically partial integro-differential equations that determine the dynamics of the “number density” of differing cell types. In PBEs, cell properties are represented as points in ℝ^*n*^, with each dimension corresponding to a different attribute. These attributes include parameters controlling cell life - for example, their rate of death and division, which vary according to a cell’s location in this “attribute” space. These functional differences control the rate at which cells progress through life, which is represented by a “flow” of cells from certain areas of attribute space to others - like chemicals diffusing down a concentration gradient. With PBEs, observed variation at a point in time is due to the initial spread of cells across attribute space coupled with the differing dynamics of cells in different areas of this space. See [11] for an introduction to PBEs.

Here, we suppose heterogeneity in quantities of interest across cells is generated by idiosyncratic variation in the rates of cellular processes. The modelling approach we follow is similar to that of [12] and is based on an ODE framework. In our model, each cell evolves according to an ODE, with its progression directed by parameters whose value varies between cells. To our knowledge, this flavour of model is unnamed, so, for sake of reference, we call them “heterogenous ODE” models (HODEs). In HODEs, the aim of inference is to estimate distributions of parameter values across cells consistent with observations. A benefit of using HODEs is that these models are computationally straightforward to simulate and, arguably, simpler to parameterise than PBEs. By using HODEs, we assume that most observed variation comes from differences in biological processes across cells, not inherent stochasticity in biochemical reactions within cells as is assumed when employing stochastic simulations algorithms.

Inference for HODEs is problematic due, partly, to the experimental hurdles involved with generating data of sufficient standard. Unlike models which represent a population by a single scalar ODE, since HODEs are individual-based, they ideally require individual cell data for estimation. A widely-used method for generating such data is flow cytometry, where a large number of cells are streamed individually through a laser beam, and, for example, the concentrations of fluorescently-labelled proteins are measured [13]. Other experimental techniques, including Western blotting and cytometric fluorescence microscopy, can also generate single cell measurements [14, 15]. These experimental methods are all, however, destructive, meaning individual cells are sacrificed during measurement, and observations at each time point hence represent *“snapshots”* of the underlying population [15]. These snapshots can be described by histograms [12] or density functions [9] fit to measurements of quantities of interest. Since HODEs assume the state of each cell evolves continuously over time, experimental data tracing individual cell trajectories through time constitutes a richer data resource. The demands of obtaining such data are, however, higher and typically involve either tracking individual cells through imaging methods [16], or trapping cells in a spatial position where they can be monitored over time [17]. These techniques impose severe restrictions on experimental practices meaning they cannot be used in many circumstances, including for online monitoring of biotechnological processes or analysis of *in vivo* studies. For this reason, “snapshot” data continues to play an important role for determining cell level variability in many applications.

By fitting HODES to snapshot data, cellular variability can be estimated and a number of approaches have been proposed for doing so. In HODEs, parameter values vary across cells according to a to-be-determined probability distribution, and the solution to the inverse problem requires solving the cell-specific ODE system many times for each individual. The count of cells in experiments typically exceeds ~ 10^4^ [15], so approaches where the computational burden scales with this count are usually infeasible. To avoid this burden, some approaches fit probability densities to raw snapshot data and use these densities, rather than raw data, for estimation [12, 15, 18, 19]. We follow this approach here. We now briefly describe the existing approaches for using HODE models to estimate cell population heterogeneity. Hasenauer et al. (2011) present a Bayesian approach to inference for HODEs, which models the input parameter space using an ansatz of a mixture of densities of chosen types. The authors then use their method to reproduce population substructure on synthetic data generated from a model of tumour necrosis factor stimulus. Hasenauer et al. (2014) use mixture models to model subpopulation structure in snapshot data with multiple-start local optimisation employed to maximise the non-convex likelihood, which they then apply to synthetic and real data from signalling pathway models. Loos et al. (2018) also use mixture models to represent subpopulation structure and use maximum likelihood to estimate both within- and between-subpopulation variability, which permits fitting to multivariate output distributions with complex correlation structures. Dixit et al. (2018) assign observations into discrete bins, then choose likelihood distributions according to the maximum entropy criterion, which they then use to estimate cell variability within a Bayesian framework.

Our framework is Bayesian although it is distinct from the approach used to fit many dynamic models, since we assume output variation arises from parameter heterogeneity across cells, with no contribution from measurement noise. The approach is, hence, most suitable when measurement error is minimal. Our method is a two-step Monte Carlo approach, which, for reasons described in §3, we call “Contour Monte Carlo” (CMC). Unlike many existing methods, CMC is straightforward to implement and does not require extensive computation time. In CMC, prior probability distributions are used in place of ansatz densities. It also does not require the number of cell clusters be chosen beforehand, rather, subpopulations emerge as modes in the posterior parameter distributions. Like [19], CMC can fit multivariate snapshot data and unlike [12], does not use discrete bins to model continuous data. As more experimental techniques elucidating single cell behaviour are developed, interest in models describing measurement snapshots should follow. We argue that due to its simplicity and generality, CMC can be used to perform inference on the proliferation of rich single cell data and, thus, is a useful addition to the modeller’s toolkit.

### Outline of the paper

In §3, we describe our probabilistic model of the inverse problem and detail the CMC algorithm for generating samples from the posterior parameter distribution. In §4, we use CMC to estimate cell population heterogeneity in three systems of biological interest.

## 3 Method

In this section, we first develop a probabilistic framework that describes our inverse problem, before introducing the CMC algorithm in pseudocode (Algorithm 1). We also detail the workflow we have found helpful in using CMC to analyse cell snapshot data (Figure 4), and suggest practical remedies to issues commonly encountered while using this approach. A glossary of variable names used in this paper is included as Table 1.

**Table 1:**
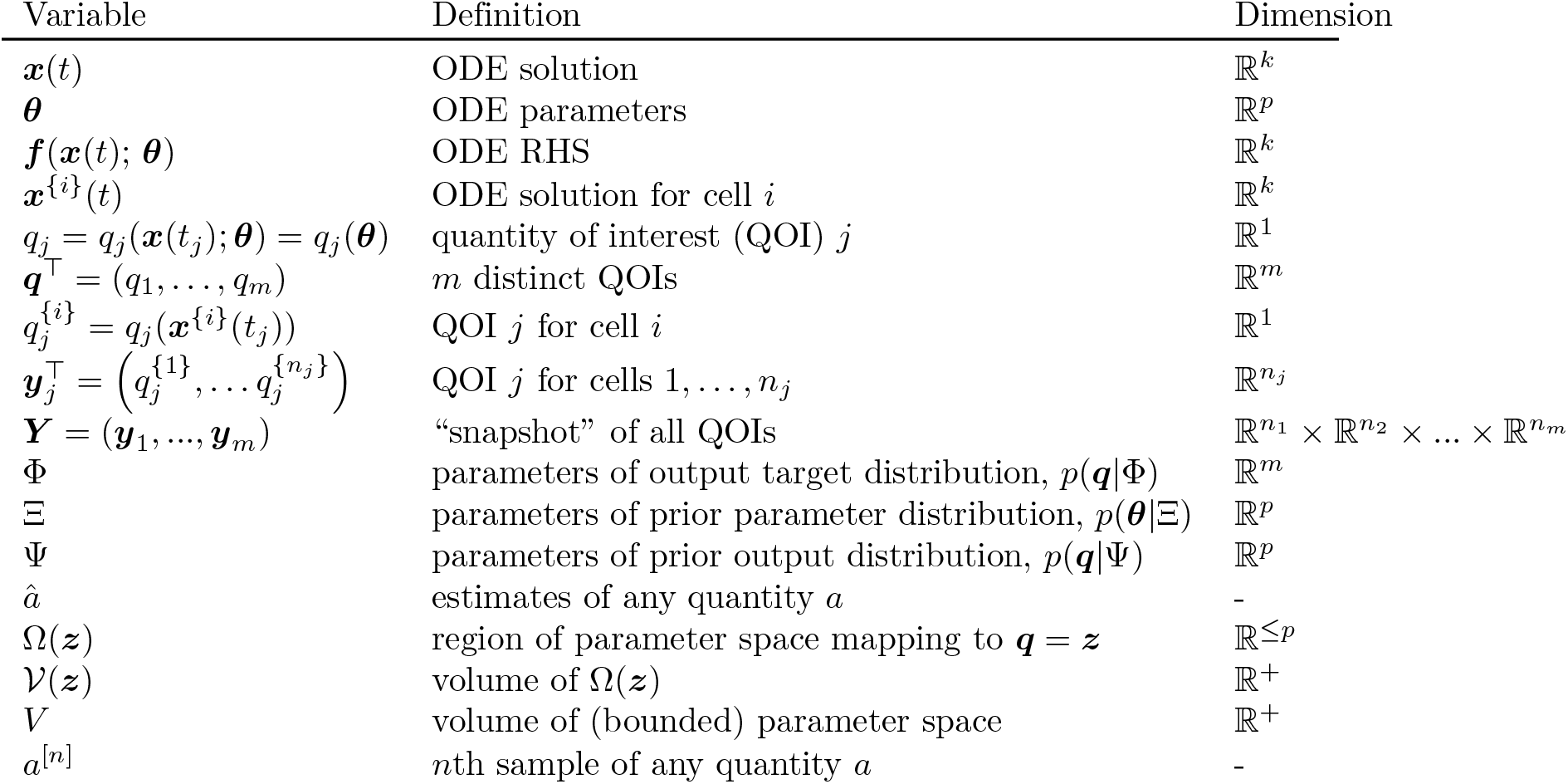
Glossary of variable names used in this paper.

Experimental methods such as flow cytometry measure single cell characteristics at a given time. Cells are typically destroyed by the measurement process, so the data consists of cross-sections or “snapshots” of sampled individuals from the population, rather than providing time series for each individual cell (Figure 1).

**Figure 1:**
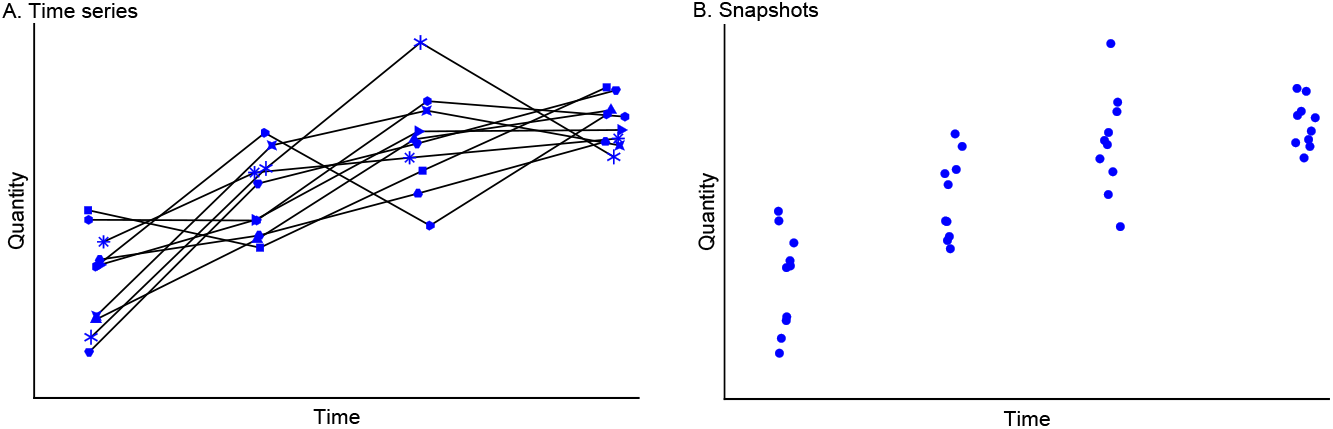
Data typical of single cell experiments. **(A) Time series data. (B) Snapshot data**. In A, note cell identities are retained at each measurement time (indicated by individual plot markers), whereas in the snapshot data in B, either this information is lost, or more often, cells are destroyed by the measurement process, and each observation corresponds to a distinct cell.

We model the processes of an individual cell using a system of ordinary differential equations (ODEs), where each element of the system typically corresponds to the concentration of a particular species. Our initial value problem is,

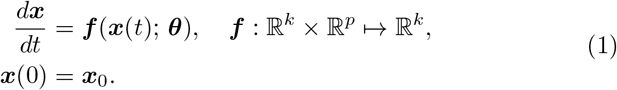

Note that in most circumstances, the initial state of the system, ***x***(0), is unknown, and it can be convenient to include these as elements of ***θ*** to be estimated.

### 3.1 Snapshot data

We assume the variation in snapshots arises due to heterogeneity in the underlying parameters, ***θ***, across cells. Therefore, the evolution of the underlying state of cell *i* is described by an idiosyncratic ODE,

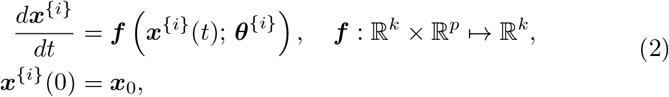

where superscript ^{*i*}^ indicates the *i*th cell. The traditional (non-hierarchical) state-space approach to modelling dynamic systems supposes that measurement error introduces stochastic variation in the output (Figure 2A). Our approach, by contrast, assumes any variation in outputs is solely due to variation in parameter values between cells (Figure 2B). Whether the assumption of “perfect” measurements is reasonable depends on experimental details of the system under investigation, but we argue our method nevertheless provides a useful approximation in cases where the signal to noise ratio is high.

**Figure 2:**
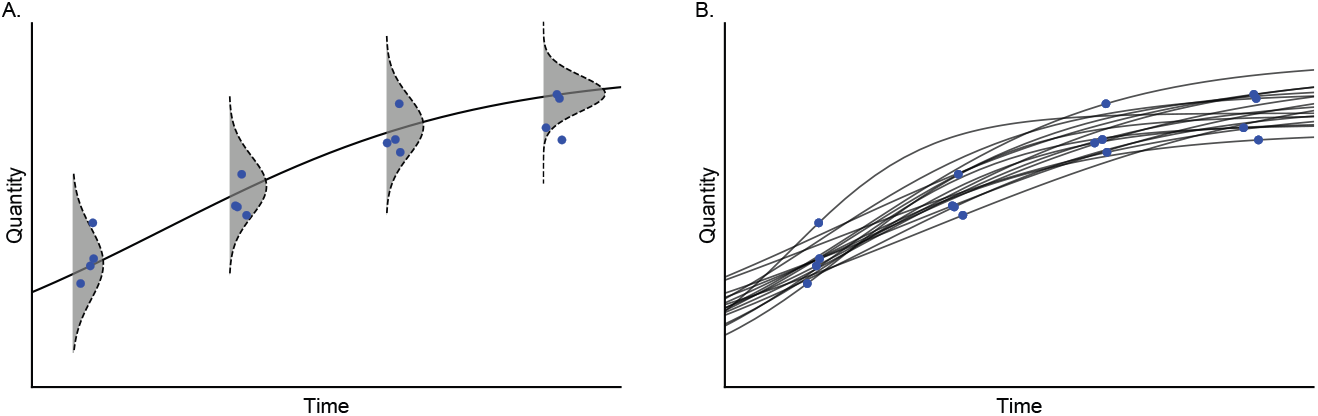
Models of variation in observed outputs. **(A) State-space model. (B) Parameter heterogeneity model**. (A) For non-hierarchical state-space models, there is a single “true” latent state, and observations result from an imperfect measurement process (grey histograms). (B) For models with parameter heterogeneity, the uncertainty is generated by differences in cellular processes (black lines) between cells. Note that, in both cases, individual cells are measured only once in their lifetime.

In an experiment, quantities of interest (QOIs) are measured. Examples of QOIs include concentrations of compounds at different points in time, peak voltages across cell membranes during an action potential, or measurements of cell volume. Here, we suppose *m* ≥ 1 QOIs are measured,

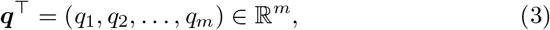

with *n*_*j*_ observations of each quantity, *q*_*j*_. Distinct QOIs, *q*_*j*_, may correspond to different functionals of the solution at the same time or the same functional at different times. The observed data for QOI *q*_*j*_ at the corresponding time *t*_*j*_ consists of the *n*_*j*_ cellular measurements,

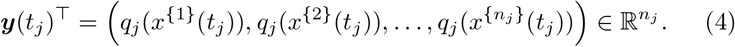

The raw snapshot data ***Y*** is the collection of all measured QOIs,

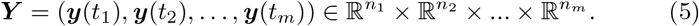

The goal of inference is to characterise the probability distribution *p*(***θ***|***Y***) representing heterogeneity in cellular processes. The numbers of cells sampled in typical experimental setups is large, and, following previous work, we represent snapshot data ***Y*** using probability distributions [12, 15, 18, 19]. In the first step of our workflow (Figure 4(i)), these distributions are approximated by a kernel density model, with support over the space of the QOI vector, ***q*** ∈ ℝ^*m*^. We use 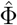 to denote the parameter estimates of the corresponding kernel density model, *p*(***q***|Φ), resultant from fitting it to raw snapshot data. We assume there are enough observational data that the estimated probability distributions are approximate sufficient statistics of the posterior distribution, meaning 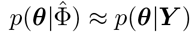.

The aim of our inverse problem, hence, becomes to derive a “posterior” parameter distribution, which, when fed through the deterministic transformation described by the model, ***q***(***θ***), recapitulates the fitted output density,

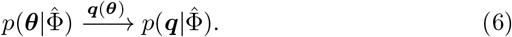

In measure theoretic terms, the intrinsic measure implied by 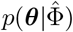 is known as the *push forward* of the measure implied by 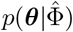 with respect to the model [20].

### 3.2 Theoretical development of CMC

We consider the under-determined case where there are fewer QOIs than model parameters (*m* < *p*). This means that, provided a given QOI can be generated by the model, it can be produced from any member of a subset of parameter space. Unlike the fully-determined case, these subsets (in general) have non-zero “volume”, and we term them “iso-output contour regions”. Symbolically, we represent the iso-output contour region for a given quantity of interest 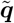 (say) by 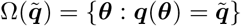.

In general, contour “volumes” 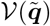 depend on the chosen output value 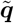 (Figure 3). Further, the interpretation of these “volumes” depends upon their dimensions. For a model with two parameters, iso-output contour regions are one-dimensional lines, whose size is a length; for a model with three parameters, contour regions are surfaces, whose size is an area; for four-dimensional parameter spaces, contour regions are three-dimensional and their size is a volume; and for models with *p* > 4 parameters, iso-contour regions are *p* − 1 dimensional manifolds, whose size is a hypervolume.

**Figure 3:**
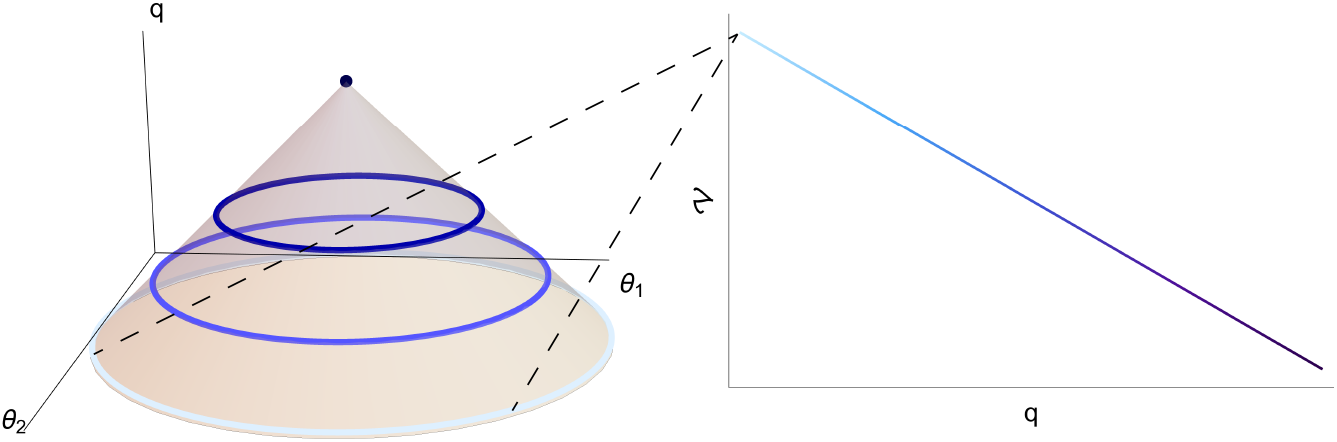
Left: An example output function *q*(*θ*_1_, *θ*_2_) along with iso-output contours indicated (coloured lines). **Right: The “volume” of output contours as a function of output value**. Note that here, since parameter space is two dimensional, the “volume” of each output value corresponds to a length of an iso-output contour.

MCMC methods aim to approximate a posterior parameter distribution by sampling from it. In this case, the resultant parameter samples, when pushed through the model, should approximate samples from the desired QOI distribution. Random Walk Metropolis [21] is a “vanilla” MCMC sampler which chooses where next to step based on the ratio of probability densities at the proposed parameter value and current position. Using a vanilla sampler for our case, unfortunately, does not work because the Markov chains are biased towards those regions of parameter space with the largest iso-output contour volumes. This bias means that the stationary parameter distribution obtained, when fed through the model, does not recapitulate the target output distribution [22].

Sampling algorithms, therefore, need to explicitly account for the differential volume of iso-output contours. In applied problems, however, we do not know the volumes of iso-output contours and they cannot be exactly calculated for all but the simplest models. Instead in CMC, we estimate them. The following analysis provides a brief introduction to a probabilistic formulation of under-determined inverse problems (see our companion paper [22] for a more comprehensive discussion). In doing so, this suggests a sampling based approach for estimating contour volumes, which are then exploited by our CMC algorithm.

Solving our inverse problem requires determining the posterior distribution of parameter values, 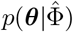, which, when used as input to the forward map, results in the target distribution, 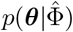. To derive the posterior parameter distribution, we consider the joint density of parameters and QOIs, 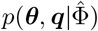. This can be decomposed in two ways using the law of conditional probability,

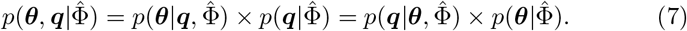

Rearranging eq. (7), we obtain the posterior parameter distribution,

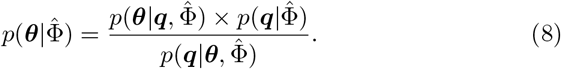

Since the mapping from parameters to outputs is deterministic, 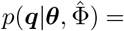 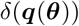, i.e., the Dirac delta function centred at ***q*** = ***q***(***θ***). Thus eq. (8) becomes,

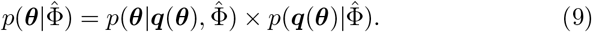

In the same way that a single output value can be caused by any member of a set of parameter values, a target output distribution 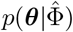 can be caused by any member of a set of parameter distributions. To ensure uniqueness of the “posterior” parameter distributions, we must, therefore, specify “prior” distributions for the parameters, as in more traditional Bayesian inference. In what follows, we assume the conditional distribution 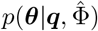 is independent of the data, i.e., 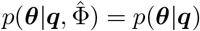, and thus represents a conditional “prior” which can be manipulated using Bayes’ rule as,

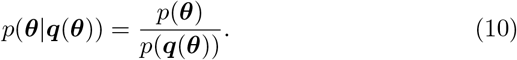

This results in the form of the posterior parameter distribution targeted by our sampling algorithm,

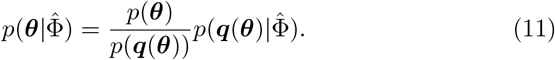

Again, we defer to our companion piece [22] for detailed explanation of eqs. (10) and (11) and, instead, here provide brief interpretation when considering a uniform prior on parameter space. In this case, 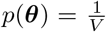, where *V* is the total volume of parameter space. The denominator term of eq. (10) is the prior induced on output space by the prior over parameter space. For a uniform prior on parameter values, this is,

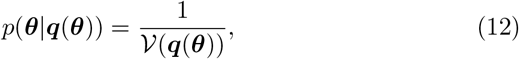

where 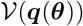 is the volume of parameter space occupied by the iso-output contour Ω(***q***(***θ***)) (see Fig. 3 for the meaning of this volume for a two parameter example). Therefore, a uniform prior over parameter space implies a prior structure where all parameter values producing the same output are given equal weighting.

### 3.3 Implementation of CMC

Except for some toy examples, the denominator of eq. (10) cannot be calculated, so exact sampling from the posterior parameter distribution of eq. (11) is not, in general, possible. We propose, instead, a computationally efficient sampling method to estimate *p*(***q***(***θ***)), which forms the first step of our so-called “Contour Monte Carlo” (CMC) algorithm (Algorithm 1; Figure 4(ii)), where the volume of iso-output contours with each feasible output value is estimated. This step involves repeated independent sampling from the prior distribution of parameters, ***θ***^[*i*]^ ~ *p*(***θ***|Ξ), where Ξ parameterises the prior probability density. Each parameter sample is then mapped to an output value, ***q***^[*i*]^ = ***q***(***θ***^[*i*]^). The collection of output samples is then fitted using a vine copula kernel density estimator (KDE) [23], 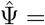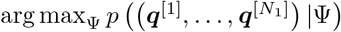. Throughout the course of development of CMC, we have tested many KDE methods and have found vine copula KDE is best suited to approximating the higher dimensional probability distributions required in practice.

The second step in our algorithm then uses MCMC to sample from an approximate version of eq. (11), where the estimated density, 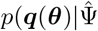 replaces its corresponding estimand (Algorithm 1; Figure 4(iii)),

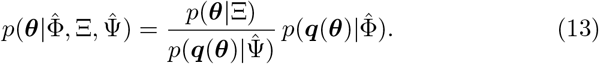

**Figure 4:**
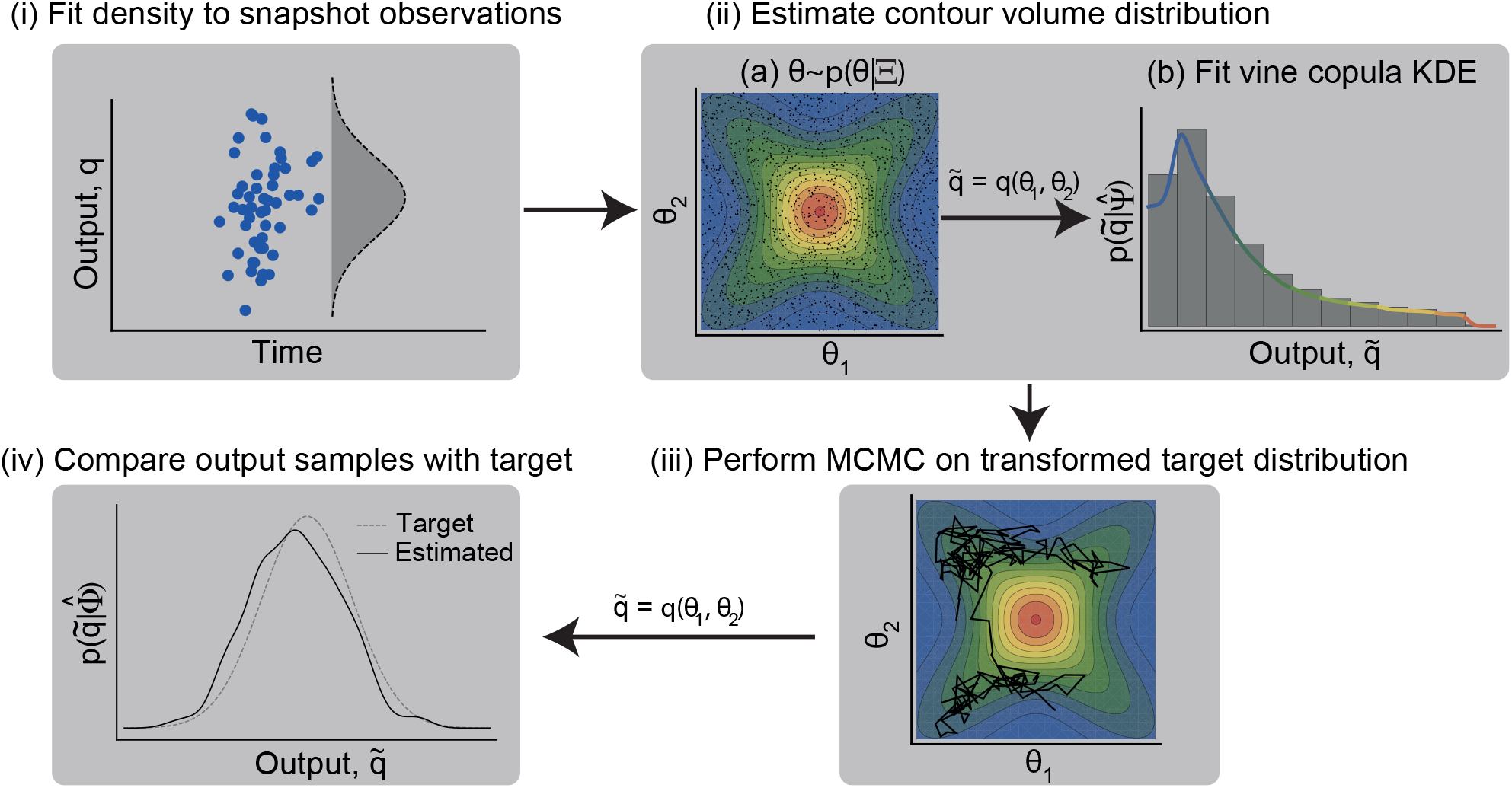
Workflow for Contour Monte Carlo to estimate cell population heterogeneity. The distribution targeted in (iii) is given by eq. (13). Here, 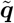 is used to represent an output value resultant from applying the functional *q* to parameter samples (*θ*_1_, *θ*_2_).

The final step in CMC is to compare output samples generated by MCMC with the target distribution (Figure 4(iv)). Asymptotically (in terms of the sample size of both sampling steps), CMC produces a sample of parameter values (***θ***^[1]^, ***θ***^[2]^,…) which, when mapped to the output space, corresponds to the target distribution 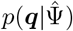. In developing CMC, we found that a finite sample of modest size for both steps of CMC results in parameter samples that, when transformed, often represented good approximations of the target. There are, however, occasions when this is not the case, and this final confirmatory step is indispensable since it frequently highlights inadequacies in contour volume estimation or MCMC, meaning more samples from either or both of these steps are required. It may also be necessary to tweak hyperparameters of the KDE in the contour volume estimation step to ensure reasonable approximation of the distribution of output values obtained by sampling the prior density. If the target distribution is sensitive to the contour volume estimates, this may also indicate that the target snapshot distribution is incompatible with the model: here, we make no claims on existence of a solution to the inverse problem, only that, Contour Monte Carlo is a pragmatic approach to approximate it by sampling if one should exist. A useful way to diagnose whether the target distribution can be produced from the model and chosen priors is to plot the output values from the contour volume estimation step of CMC - this is akin to visualising the prior predictive distribution in traditional Bayesian inference [21]. If the bulk of target probability mass does not overlap with the simulated output values, then the model and/or chosen prior are unlikely to be invertible to this particular target.

### 3.4 Workflow and CMC algorithm

A graphical illustration of the complete CMC workflow is provided in Figure 4. All variables are defined in Table 1. The CMC algorithm is provided in Algorithm 1. In this implementation, MCMC sampling is performed via the Random Walk Metropolis algorithm, but for the examples in §4, we use an adaptive MCMC algorithm [24].

**Algorithm 1.**
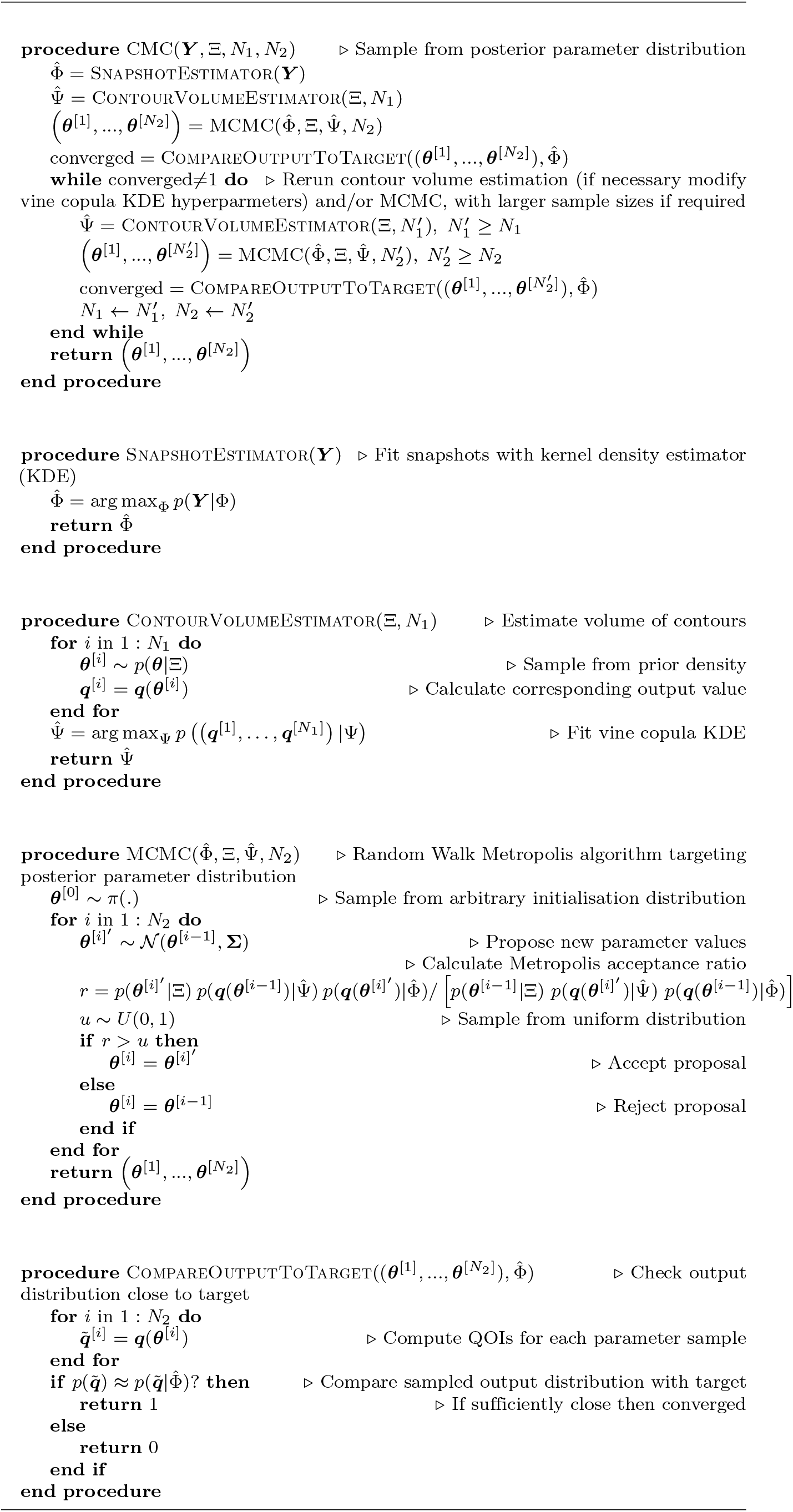
Pseudocode for the Contour Monte Carlo algorithm for sampling from the posterior parameter distribution of eq. (13).

To generate our results in §4, we assumed for the contour volume estimation step sample sizes were sufficient if the output samples from MCMC provided a reasonable approximation to the target, although we recognise that future work should refine this process further. For the MCMC step, we used adaptive covariance MCMC (see SOM of [24]) to sample from the target distribution, as it typically provides a considerable speed-up over Random Walk Metropolis [21, 25]. We also used the Gelman-Rubin convergence statistic, 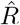, to diagnose convergence [21,26], with a convergence threshold of 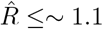.

To solve the forward model of each differential equation, we used Julia’s inbuilt “solve” method for ODE models, which automatically chooses an efficient inbuilt solver [27]. To replicate the results in this section, we recommend readers execute the corresponding Julia scripts (one for each result section) at https://github.com/ben18785/inverse-sensitivity/tree/master/examples. Note that, these scripts use the “*RCall* ” library for Julia [28], which calls `R` from Julia. This package was necessary to use the “*kdevine*” `R` package for vine copula kernel density estimation [29].

## 4 Results

In this section, we use CMC to estimate posterior parameter distributions for three biological systems. In all but one of the examples, we assume that the first step of CMC (“SnapshotEstimator” within Algorithm 1) has already been completed, and we are faced with inferring a parameter distribution which, when mapped to outputs, recapitulates the target density. To accompany the text, we provide the Julia notebook used to generate the results. A table of priors used for each example is provided in Table 3.

### 4.1 Growth factor model

We first consider the “growth factor model” introduced by [12], which concerns the dynamics of inactive ligand-free cell surface receptors, *R*, and active ligand-bound cell surface receptors, *P*, modulated by an exogenous ligand, *L*. The governing dynamics are determined by the following system,

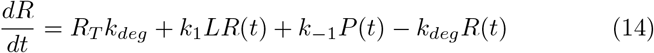

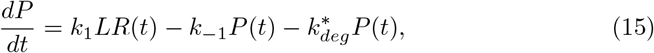

with initial conditions,

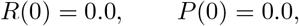

where 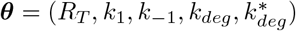 are parameters to be determined. In this example, we use measurements of the active ligand-bound receptors *P* to estimate cellular heterogeneity in these processes. We denote the solution of eq. (15) as *P* (*t*; ***θ***, ***L***) and seek to determine the parameter distribution consistent with an output distribution,

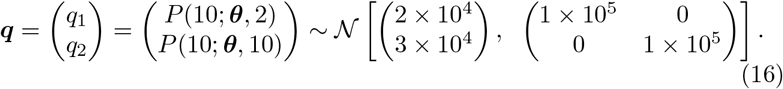

#### 4.1.1 Uniform prior

To start, we specify a uniform prior for each of the five parameters, with bounds given in Table 3, and use CMC to estimate the posterior parameter distribution. In Figure 5A, we show the sampled outputs (blue points) versus the contours of the target distribution (black solid closed curves), illustrating a good correspondence between the sampled and target densities. Above and to the right of the main panel, we also display the marginal target densities (solid black lines) versus kernel density estimator reconstructions of the output marginals from the CMC samples (dashed blue lines), which again highlights the fidelity of the CMC sampled density to the target.

**Figure 5:**
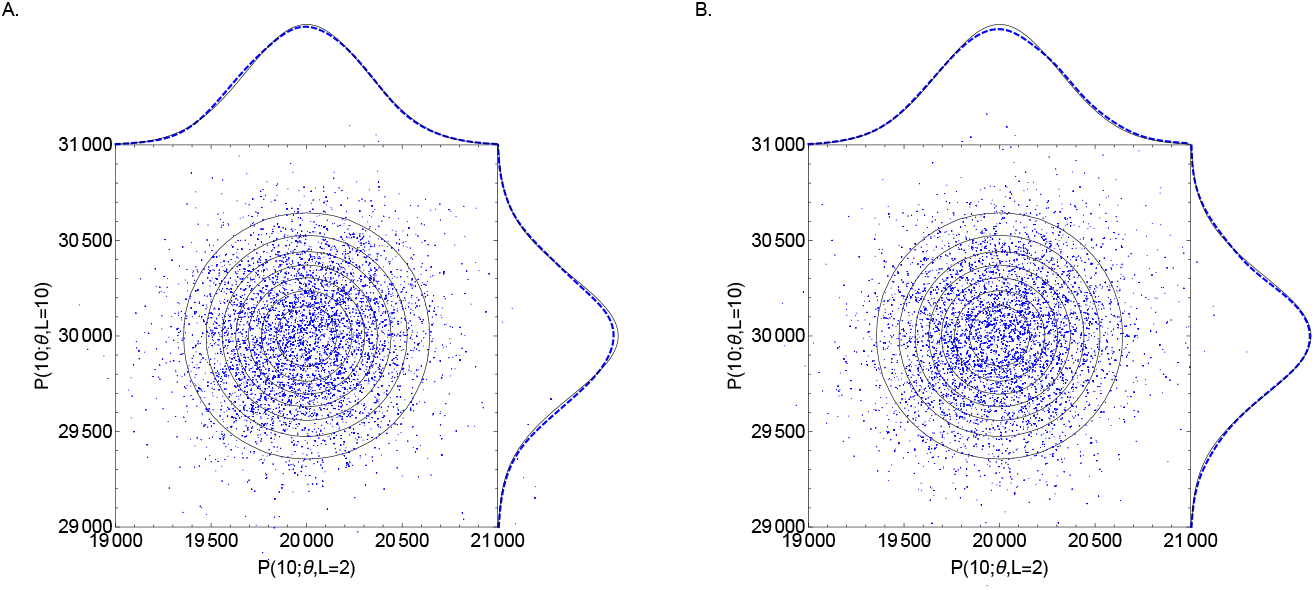
Growth factor model. Target joint output distribution (solid contour lines) and target marginal distributions (solid lines; above and to the right of each figure) versus outputs sampled by CMC (blue points) and reconstructed marginals (dashed lines). (A) uniform priors. (B) Gaussian priors. In CMC, 100,000 independent samples were used in the “ContourVolumeEstimator” step and 10,000 MCMC samples across each of 4 Markov chains were used in the second step, with the first half of the chains discarded as “warm-up” [21]. For the reconstructed marginal densities in the plots, we use Mathematica’s “SmoothKernelDistribution” function specifying bandwidths of 100 with Gaussian kernels [30].

In Figure 6A, we plot the joint posterior parameter distribution for *k*_1_, the rate of ligand binding to inactive receptors and *k*_−1_, which dictates the rate of the reverse reaction. A given level of bound ligands can be generated in many different ways. Not surprisingly, it is the *ratio* of the forward and reverse reaction rates, *k*_1_ and *k*_−1_ respectively, that is of greatest importance, and because of this, the distribution representing cell process heterogeneity contains linear positive correlations between these parameters.

**Figure 6:**
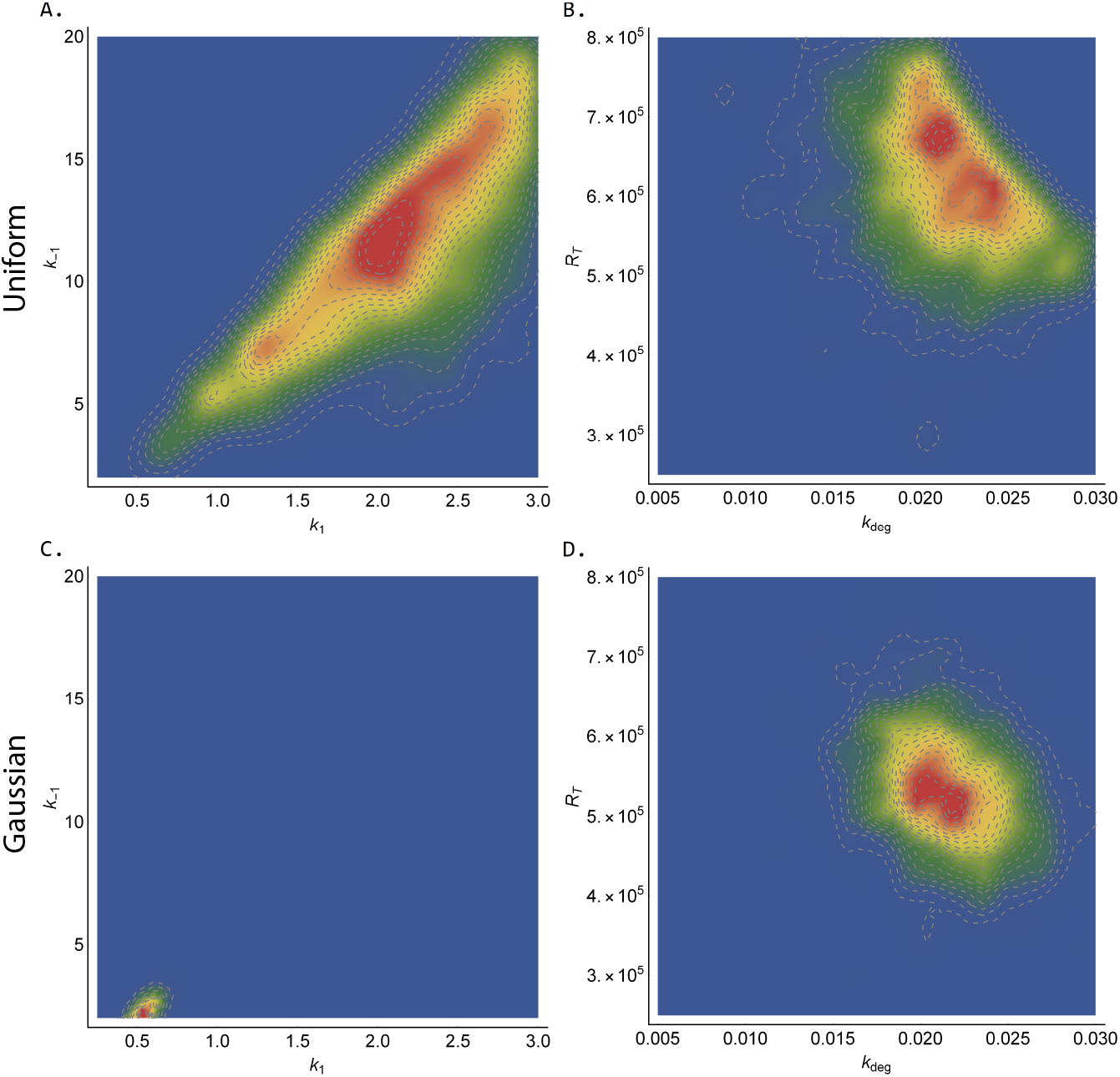
Growth factor model. Joint posterior distributions estimated by CMC. Top row (A-B): (*k*_1_, *k*_−1_) and(*k*_*deg*_, *R*_*T*_) using uniform priors. Bottom row (C-D):(*k*_1_, *k*_−1_) and(*k*_*deg*_, *R*_*T*_) using Gaussian priors. See Figure 5 caption for CMC details and Table 3 for the priors used. Red (blue) indicates areas of relatively high (low) probability density.

In Figure 6B, we show the posterior parameter distribution for *k*_*deg*_, the rate of degradation of ligand-free cell surface receptors and *R*_*T*_, the rate of introduction of ligand-free cell surface receptors. This plot shows more concentrated posterior mass than in Figure 6A. Why do our measurements allow us to better resolve (*k*_*deg*_, *R*_*T*_) compared to (*k*_1_, *k*_−1_)? To answer this, it is useful to calculate the sensitivity of *P* (*t*; ***θ***, ***L***) to changes in each of the parameters. To account for the differing magnitudes of each parameter, we calculate elasticities, the proportional changes in measured output for a proportional change in parameter values, using the forward sensitivities method described in [31], and these are shown in Figure 7. When the exogenous ligand is set at *L* = 2, these indicate the active ligand-bound receptor concentration is most elastic to changes in *R*_*T*_ and *k*_*deg*_. This higher elasticity means that their range is more restricted by the output measurement than for *k*_1_ and *k*_−1_, which have much smaller elasticities at *t* = 10. In Table 2, we show the posterior quantiles for the estimated parameters, and in the last column, indicate the ratio of the 25%-75% posterior interval widths to the uniform prior range for each parameter. These were strongly negatively correlated with the magnitude of the elasticities for each parameter (*ρ* = 0.95, *t* = −5.22, *df* = 3, *p* = 0.01 for Pearson’s product-moment correlation), indicating the utility of sensitivity analyses for optimal experimental design. We suggest, however, that CMC can also be used for this purpose. If an experimenter generates synthetic data for various choices of QOIs, they can use CMC to derive the posterior parameter distributions in each case. They then, simply, select the particular QOI producing the narrowest posterior for key parameters.

**Table 2:** Growth factor model. Estimated quantiles from CMC samples with uniform and Gaussian priors. The last column indicates the proportion of the uniform prior bounds occupied by the 25%-75% posterior interval in each case. The prior hyperparameters used in each case are given in Table 3.

| Parameter | Quantiles |  |  |  |  | Posterior<br>25%-75%<br>conc. |
| --- | --- | --- | --- | --- | --- | --- |
|  | 2.5% | 25% | 50% | 75% | 97.5% |  |
| Uniform prior |  |  |  |  |  |  |
| $R_T$ | 441,006 | 548,275 | 606,439 | 677,055 | 772,484 | 23% |
| $k_1$ | 0.90 | 1.69 | 2.17 | 2.56 | 2.95 | 32% |
| $k_{-1}$ | 4.35 | 8.35 | 11.23 | 14.23 | 18.71 | 33% |
| $k_{deg}$ | 0.013 | 0.019 | 0.021 | 0.024 | 0.029 | 20% |
| $k_{deg}^*$ | 0.20 | 0.34 | 0.40 | 0.44 | 0.49 | 27% |
| Gaussian prior |  |  |  |  |  |  |
| $R_T$ | 408,396 | 487,372 | 529,558 | 577,970 | 678,632 | 16% |
| $k_1$ | 0.39 | 0.49 | 0.54 | 0.60 | 0.70 | 4% |
| $k_{-1}$ | 1.39 | 1.92 | 2.26 | 2.63 | 3.35 | 4% |
| $k_{deg}$ | 0.016 | 0.020 | 0.022 | 0.024 | 0.027 | 16% |
| $k_{deg}^*$ | 0.22 | 0.29 | 0.33 | 0.38 | 0.46 | 21% |

**Table 3:** Priors used for each example in §4. The parameters *p*_1_ and *p*_2_ indicate the prior hyperparameters: for uniform priors, these correspond to the lower and upper limits; for Gaussian priors, they correspond to the mean and standard deviation.

| Model | Target density | Parameter | Prior density | Prior $p_1$ | Prior $p_2$ |
| --- | --- | --- | --- | --- | --- |
| Growth factor | 2D Gaussian | $R_T$ | uniform | $2.5 \times 10^5$ | $8 \times 10^5$ |
| | | $k_1$ | uniform | 0.25 | 3.0 |
| | | $k_{-1}$ | uniform | 2.0 | 20.0 |
| | | $k_{deg}$ | uniform | 0.005 | 0.03 |
| | | $k_{deg}^*$ | uniform | 0.1 | 0.5 |
| Growth factor | 2D Gaussian | $R_T$ | Gaussian | $5 \times 10^5$ | $1 \times 10^5$ |
| | | $k_1$ | Gaussian | 0.5 | 0.1 |
| | | $k_{-1}$ | Gaussian | 3.0 | 1.0 |
| | | $k_{deg}$ | Gaussian | 0.02 | 0.005 |
| | | $k_{deg}^*$ | Gaussian | 0.3 | 0.1 |
| Michaelis-Menten | bimodal Gaussian | $k_f$ | uniform | 0.2 | 15 |
| | | $k_r$ | uniform | 0.2 | 2.0 |
| | | $k_{cat}$ | uniform | 0.5 | 3.0 |
| Michaelis-Menten | 4D Gaussian | $k_f$ | uniform | 0.2 | 15 |
| | | $k_r$ | uniform | 0.2 | 2.0 |
| | | $k_{cat}$ | uniform | 0.2 | 3.0 |
| | | $E_0$ | uniform | 3.0 | 5.0 |
| | | $S_0$ | uniform | 5.0 | 10.0 |
| | | $C_0$ | uniform | 0.0 | 0.2 |
| | | $P_0$ | uniform | 0.0 | 0.2 |
| TNF signalling | bivariate Gaussian | $a_1$ | uniform | 0.4 | 0.8 |
| | | $a_2$ | uniform | 0.1 | 0.7 |
| | | $a_3$ | uniform | 0.3 | 0.7 |
| | | $a_4$ | uniform | 0.1 | 0.3 |
| | | $b_1$ | uniform | 0.5 | 0.7 |
| | | $b_2$ | uniform | 0.4 | 0.6 |
| | | $b_3$ | uniform | 0.4 | 0.6 |
| | | $b_4$ | uniform | 0.2 | 0.4 |
| | | $b_5$ | uniform | 0.2 | 0.4 |
| TNF signalling | bimodal Gaussian | $a_1$ | uniform | 0.5 | 0.7 |
| | | $a_2$ | uniform | 0.1 | 0.3 |
| | | $a_3$ | uniform | 0.1 | 0.3 |
| | | $a_4$ | uniform | 0.4 | 0.6 |
| | | $b_1$ | uniform | 0.3 | 0.5 |
| | | $b_2$ | uniform | 0.6 | 0.8 |
| | | $b_3$ | uniform | 0.2 | 0.4 |
| | | $b_4$ | uniform | 0.4 | 0.6 |
| | | $b_5$ | uniform | 0.3 | 0.5 |

**Figure 7:**
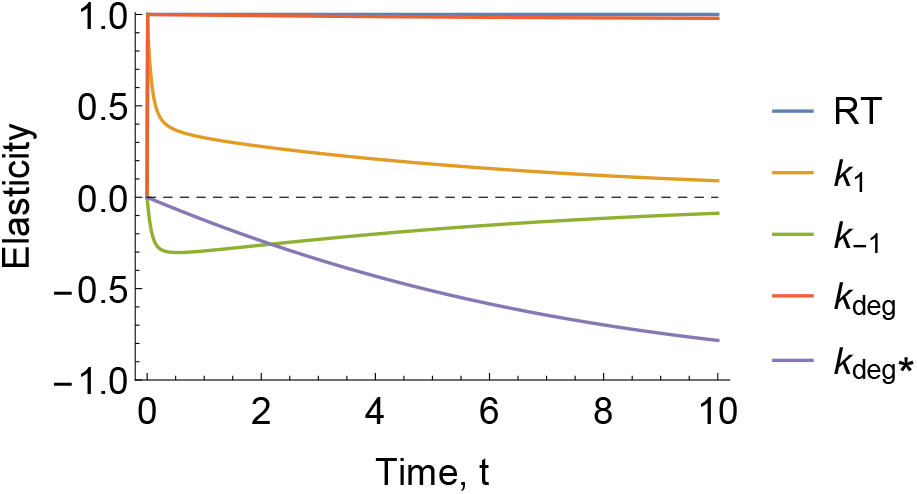
Growth factor model. **Elasticities of the active ligand-bound receptors *P* with respect to each parameter as a function of time**. When calculating the elasticities of each parameter, the other parameters were set to their posterior medians given in Table 2 and *L* = 2.

#### 4.1.2 Gaussian prior

For an under-determined model, the number of QOIs, *m*, is less than the number of parameters, *p*, and there typically exists a non-singular set of parameter distributions mapping to the same target output distribution. To uniquely identify a posterior parameter distribution, it is, therefore, necessary to specify a prior parameter distribution. By incorporating priors, this allows pre-existing biological knowledge to be included, leading to reduced uncertainty in parameter estimates. CMC allows any prior with correct support to be used. Changes to priors affect both the “ContourVolumeEstimation” and “MCMC” steps of CMC (Algorithm 1), so that the (changed) posterior parameter distribution still maps to the target.

We now use CMC to estimate the posterior parameter distribution, when using Gaussian priors (prior hyperparameters shown in Table 3), which are more concentrated than the uniform priors used in §4.1.1. As desired, the target output distribution appears virtually unaffected by the change of priors (Figure 5B) although with substantial changes to the posterior parameter distribution (Figure 6C and 6D). In particular, the marginal posterior distributions obtained from the Gaussian prior are narrower compared to the uniform case (rightmost column of Table 2).

As in traditional Bayesian inference, prior choice has a greater influence on the posterior distribution when data provide less information on the underlying process. This is readily apparent in comparing the dramatic change from Figure 5A to 5C for (*k*_1_, *k*_−1_), which have low sensitivities, with the more nuanced change from Figure 5B to 5D for (*k*_*deg*_, *R*_*T*_), which have high sensitivities.

### 4.2 Michaelis-Menten kinetics

In this section, we use CMC to invert output measurements from the Michaelis-Menten model of enzyme kinetics (see, for example, [32]) `- il-` lustrating how CMC can determine resolve population substructure from a multimodal output distribution. The Michaelis-Menten model of enzyme kinetics describes the dynamics of concentrations of an enzyme, *E*, a substrate, *S*, an enzyme-substrate complex, *C*, and a product, *P*,

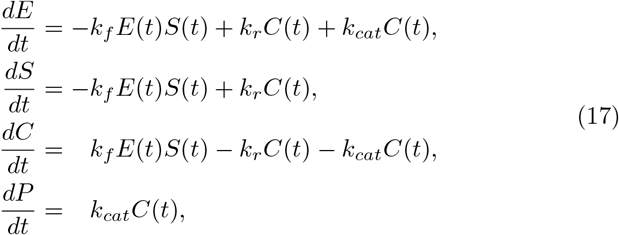

with initial conditions,

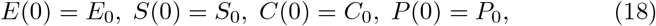

where *k*_*f*_ is the rate of the forward reaction *E* + *S* → *C*, *k*_*r*_ is the rate of the reverse reaction *C* → *E* + *S*, and *k_cat_* is the catalytic rate of product formation by the reaction *C* → *E* + *P* .

#### 4.2.1 Bimodal output distribution

When subpopulations of cells, each with distinct dynamics, are thought to exist, determining their characteristics - the proportions of cells in each cluster, their distinct parameter values, and so on - is often of key interest [15, 19]. Before formal inference occurs, an output distribution with multiple modes may signal the existence of fragmented subpopulations of cells, and to exemplify this, we target a bimodal bivariate Gaussian distribution for measurements of the level of enzyme and substrate at *t* = 1 and *t* = 2 respectively,

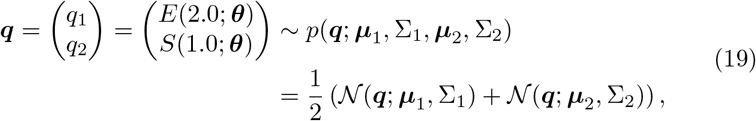

where ***θ*** = (*k*_*f*_, *k*_*r*_, *k*_*cat*_). The parameters of the Gaussian mixture components are,

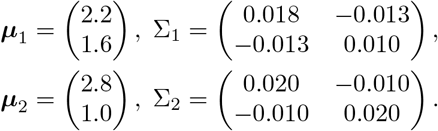

In what follows, we specify uniform priors on each element of ***θ*** (see Table 3). Using a modest number of samples in each step, CMC provides a close approximation to the output target distribution (Figure 8A). Without providing *a priori* information on the subpopulations of cells, two distinct clusters of cells emerged from application of CMC (orange and blue points in Figure 8B) - each corresponding to distinct modes of the output distribution (corresponding coloured points in Figure 8A). It is worth noting, however, that the issues inherent with using MCMC to sample multimodal distributions similarly apply here. So, whilst adaptive MCMC [24] sufficed to explore this posterior surface, it may be necessary to use MCMC methods more robust to such geometries in other cases (for example, population MCMC [33]).

**Figure 8:**
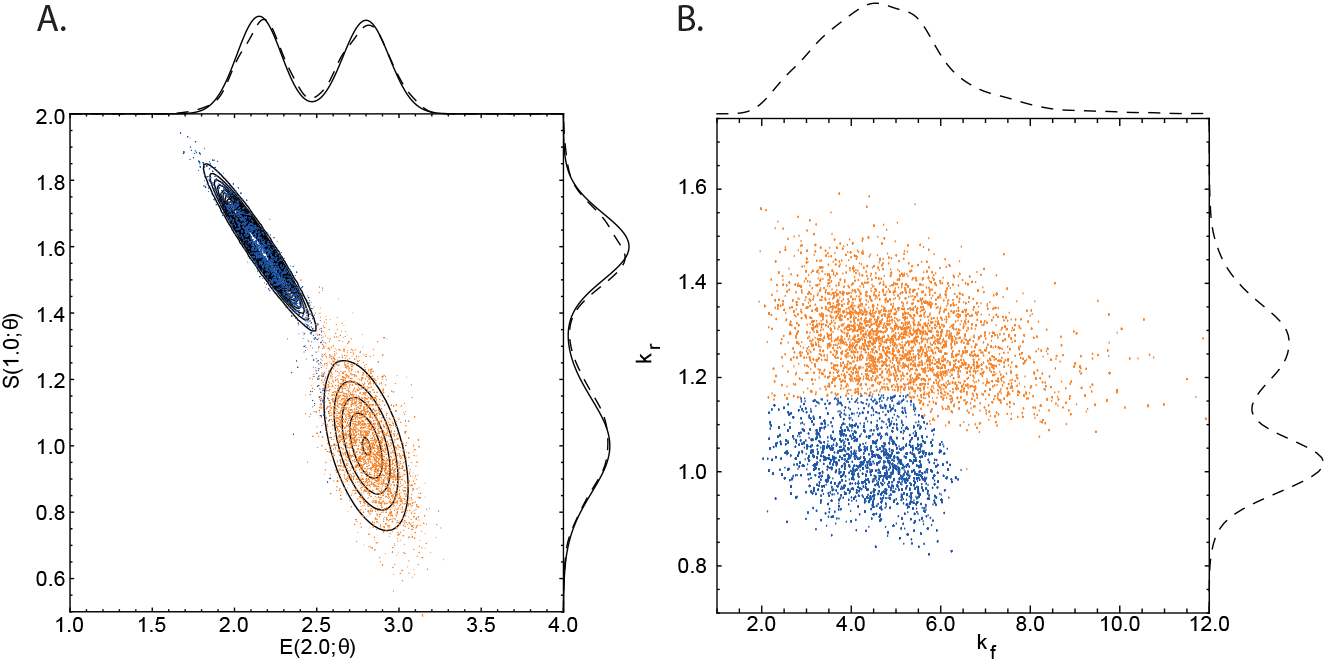
Michaelis-Menten model. **(A) Bimodal target distribution *q* (solid contour lines) versus output samples (points). (B) posterior parameter samples (points)**. The solid and dashed lines above and to the side of panel A indicate the target and estimated marginal output distributions, respectively. In B, only estimated parameter marginals are shown as the exact solutions are unknown. The orange (blue) points in A were generated by the orange (blue) parameter samples in B. See Figure 5 caption for CMC details. Mathematica’s “SmoothKernelDistribution” function [30] with Gaussian kernels was used to construct marginal densities with: (A) default bandwidths, and (B) bandwidths of 0.3 (horizontal axis) and 0.03 (vertical axis). Mathematica’s “ClusteringComponents” function [30] was used to identify clusters in B.

#### 4.2.2 Four-dimensional output distribution

Loos et al. (2018) consider a multidimensional output distribution, with correlations between system characteristics that evolve over time. Our approach allows arbitrary covariance structure between measurements, and to exemplify this, we now target a four-dimensional output distribution, with paired measurements of enzyme and substrate at *t* = 1 and *t* = 2,

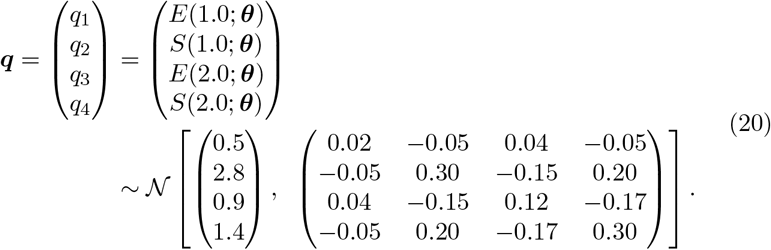

Since this target has four QOIs, and the Michaelis-Menten model has three rate parameters (*k*_*f*_, *k*_*r*_, *k*_*cat*_), the system is over-identified and so CMC cannot be straightforwardly applied. Instead, we allow the four initial states (*E*_0_, *S*_0_, *C*_0_, *P*_0_) to be uncertain quantities, bringing the total number of parameters to seven. We set uniform priors on all parameters (see Table 3). In order to check that the model and priors were consistent with the output distribution given by eq. (20), we plotted the output measurements used to estimate contour volumes (obtained from the first step of the “ContourVolumeEstimator” method in Algorithm 1) against the target (Figure 9). Since the main support of the densities (black contours) lies within a region of output space reached by independent sampling of the priors (blue points), this indicated the target could feasibly be generated from this model and priors, and we proceeded to estimation by CMC.

**Figure 9:**
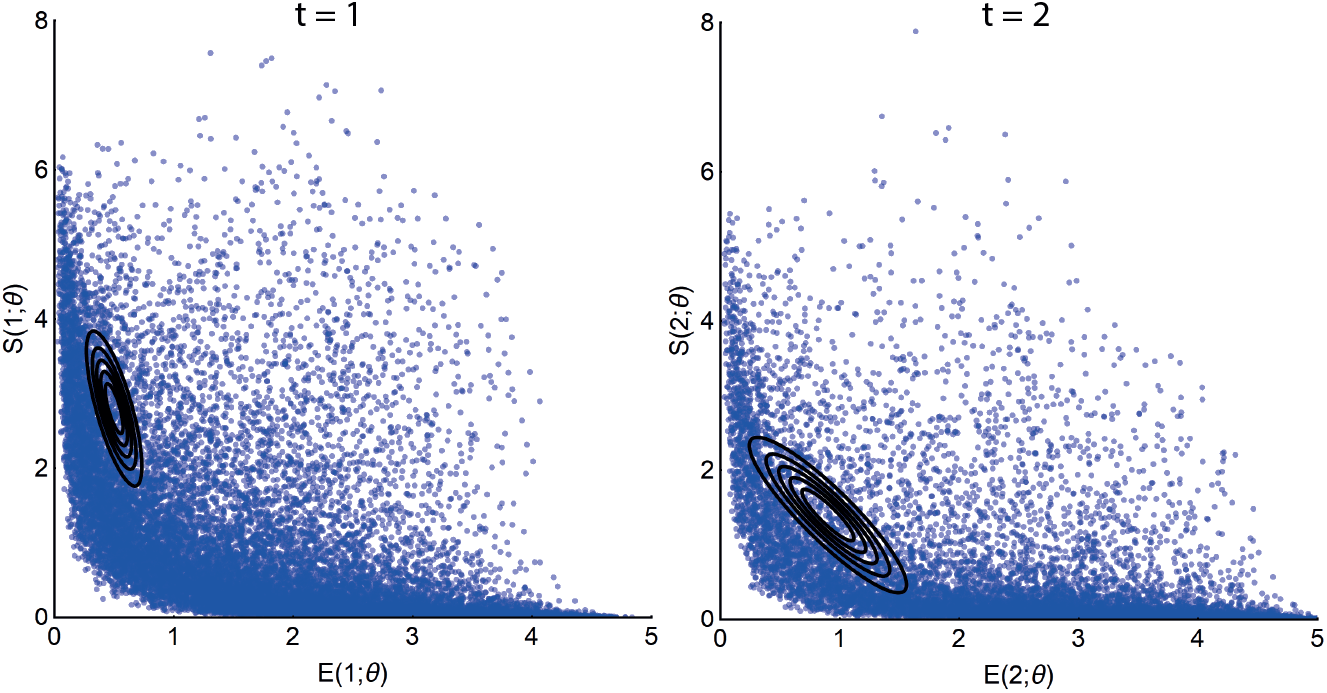
Michaelis-Menten model. **QOIs (blue points) obtained by independently sampling the priors versus the target distribution (black solid contours). Left: (*q*_1_, *q*_2_). Right:(*q*_3_, *q*_4_)**. We show 20,000 output samples, where each set of four measurements was obtained from a single sample of all parameters. The output target distribution shown by the contours corresponds to the marginal densities of each pair of enzyme-substrate measurements given by eq. (20).

Figure 10 plots the output samples of enzyme and substrate from the last step of CMC for *t* = 1 (blue points) and *t* = 2 (orange points) versus the contours (black lines) of the joint marginal distributions of eq. (20). The distribution of paired enzyme-substrate samples illustrates that the CMC output distribution closely approximates the target density, itself representing dynamic evolution of the covariance between enzyme and substrate measurements. Target marginal distributions (solid lines) along with their approximations from kernel density estimation (dashed lines) are also shown above and to the right of the main panel of Figure 10 and largely indicate correspondence.

**Figure 10:**
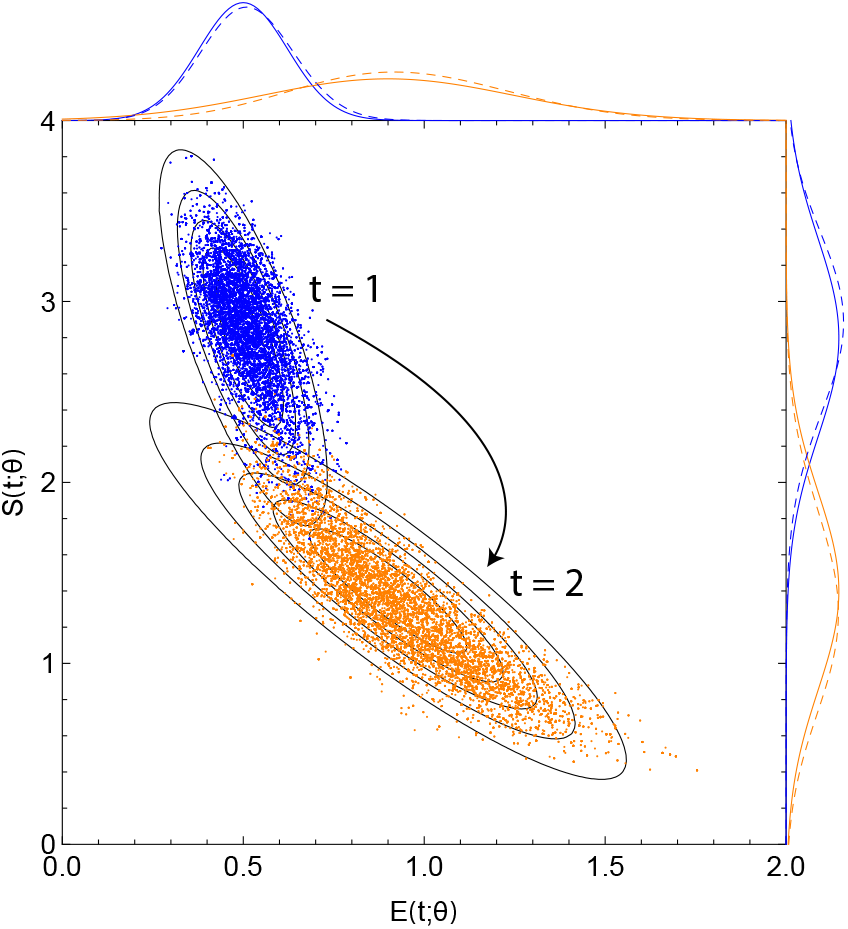
Michaelis-Menten model. **Posterior output samples from CMC (coloured points) versus contour plots (black solid lines) of the joint marginal distributions of eq. (20)**. Enzyme and substrate measurements are given by the horizontal and vertical axes, respectively. Output functionals for (*q*_1_, *q*_2_) and (*q*_3_, *q*_4_) are given by blue and orange points, respectively. The solid and dashed coloured lines outside the panels indicate exact target marginals of eq. (20) and those estimated by CMC, respectively. In the “ContourVolumeEstimator” step, 200,000 independent samples were used, and in the MCMC step, 10,000 samples across each of 4 Markov chains were used, with the first half of the chains discarded as “warm-up” [21]. Mathematica’s “SmoothKernelDistribution” function, using Gaussian kernels [30] and bandwidths ranging from 0.1 to 0.4, was used to reconstruct marginal densities.

### 4.3 TNF signalling pathway

We now illustrate how CMC can be applied to an ODE system of larger size: the tumour necrosis factor (TNF) signalling pathway model introduced in [34] and used by [15] to illustrate a Bayesian approach to cell population variability estimation. The model incorporates known activating and inhibitory interactions between four key species within the TNF pathway: active caspase 8, *x*_1_, active caspase 3, *x*_2_, a nuclear transcription factor, *x*_3_ and its inhibitor, *x*_4_, such that

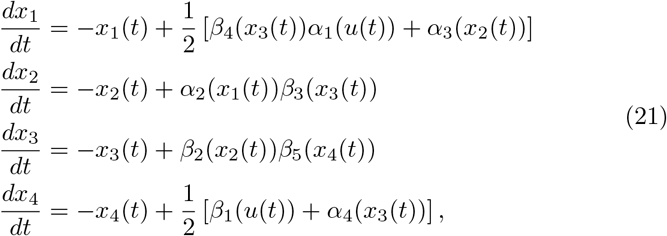

with initial conditions,

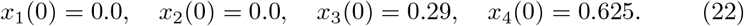

The functions *α*_*i*_ and *β*_*j*_ represent activating and inhibitory interactions respectively,

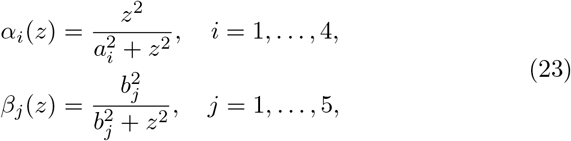

and the parameters *a*_*i*_ for *i* ∈ (1, 2, 3, 4) and *b*_*j*_ for *j* ∈ (1, 2, 3, 4, 5) represent activation and inhibition thresholds. The function *u*(*t*) represents a TNF stimulus represented by a top hat function,

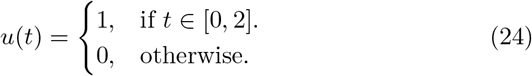

#### 4.3.1 Recovering parameter values in under-determined systems

In under-determined models, a set of parameters of non-zero volume can produce the same output values. A consequence of this unidentifiability is that we cannot perform “full circle” inference: that is, using a known parameter distribution to generate an output distribution does not result in that parameter distribution being recovered through inference. We illustrate this idea by generating an output distribution by varying a single parameter value between runs of the forward model (21) and performing inference on all nine system parameters, whilst collecting only two output measurements. Specifically, we randomly sample 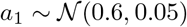 for each simulation of the forward model, whilst holding the other parameters constant,

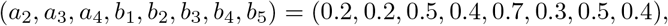

and measure *q*_1_ = *x*_1_(2.0) and *q*_2_ = *x*_2_(1.0) in each case. In doing so, we obtain an output distribution well-approximated by the bivariate Gaussian distribution,

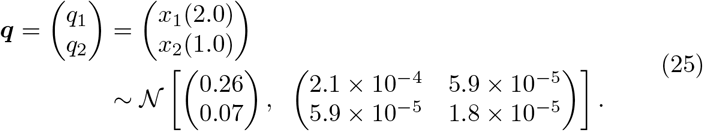

We now apply CMC to the target output distribution given by eq. (25) to estimate a posterior distribution over all nine parameters of eq. (21). Apart from a few cases, the priors for each parameter were chosen to *exclude* the values that were used to generate the output distribution (see Table 3), to illustrate how the recovered posterior distribution and data generating distribution differ. In Figure 11A, we plot the actual parameter values (horizontal axis) used to generate the data versus the estimated values (vertical axis). This illustrates that, due to the chosen priors, there is a disjunction between actual and estimated parameter values in all cases apart from *a*_1_. Though because the model is under-determined, the corresponding output distribution closely approximates the target despite these differences (Figure 11B).

**Figure 11:**
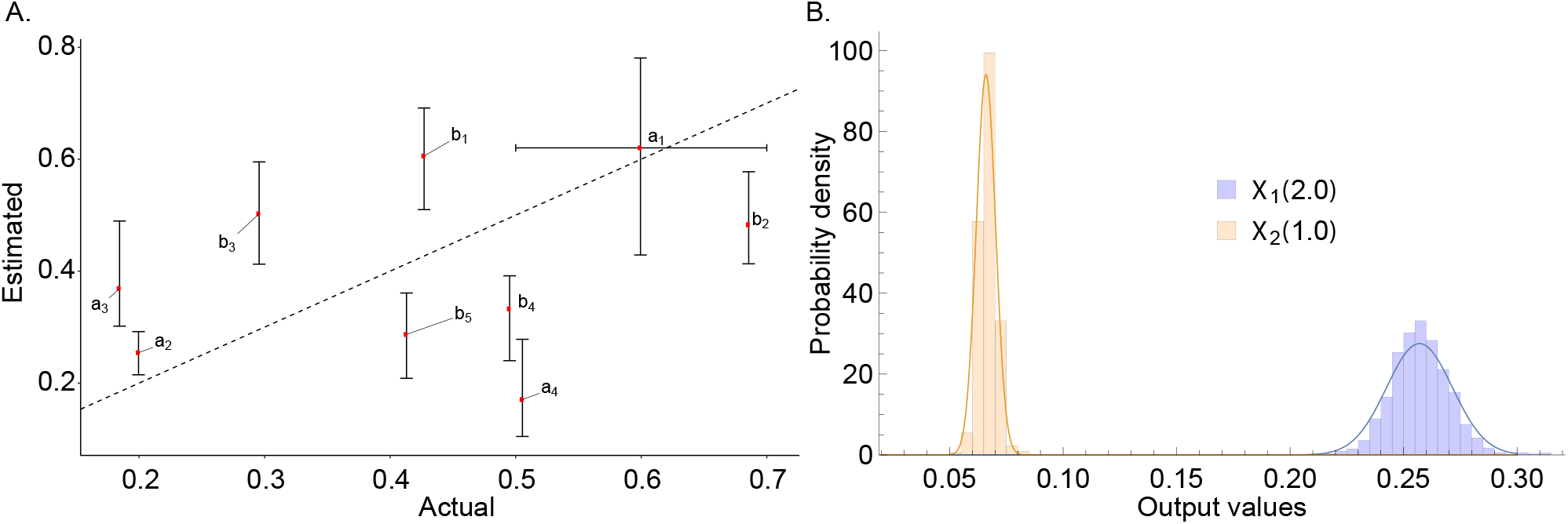
TNF signalling pathway model. (A) Actual parameter values versus estimated quantiles for the output distribution of eq. (25). (B) Marginal output targets (solid lines) and sampled output distributions (histograms). In A, in the vertical direction, red points indicate 50% posterior quantiles and upper and lower whiskers indicate 97.5% and 2.5% quantiles, respectively; in the horizontal direction, with the exception of *a*_1_, red points indicate the parameter values used to generate the data; for *a*_1_, the red point indicates the mean of the Gaussian distribution used to generate the data and the whiskers indicate its 95% quantiles. In CMC, 10,000 independent samples were used in the “ContourVolumeEstimator” step, and 5,000 MCMC samples across each of 4 Markov chains were used in the second, with the first half of the chains discarded as “warm-up” [21].

#### 4.3.2 Bimodal output distribution

The dynamics of all cells can often be modelled by assuming cells exist in subpopulation clusters, which evolve differently over time. A hint that such subpopulation structure may exist is output distributions with multiple modes. We now apply CMC to investigate a bimodal output distribution for the TNF signalling pathway model similar to that investigated by [15]. We aim to estimate the posterior parameter distribution mapping to the following output distribution,

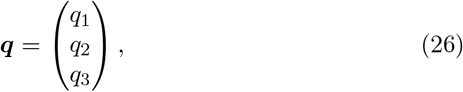

where,

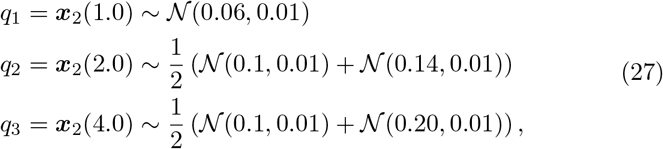

where the target distributions for ***q***_2_(2.0) and ***q***_2_(4.0) indicate mixtures of univariate Gaussians, and the priors used are given in Table 3. This target distribution, along with the unique trajectories obtained by applying the CMC algorithm, are shown in Figure 12. This figure illustrates that a bimodal output distribution causes CMC to sample clusters of parameter values, without the need for subpopulation information to be provided ahead of estimation.

**Figure 12:**
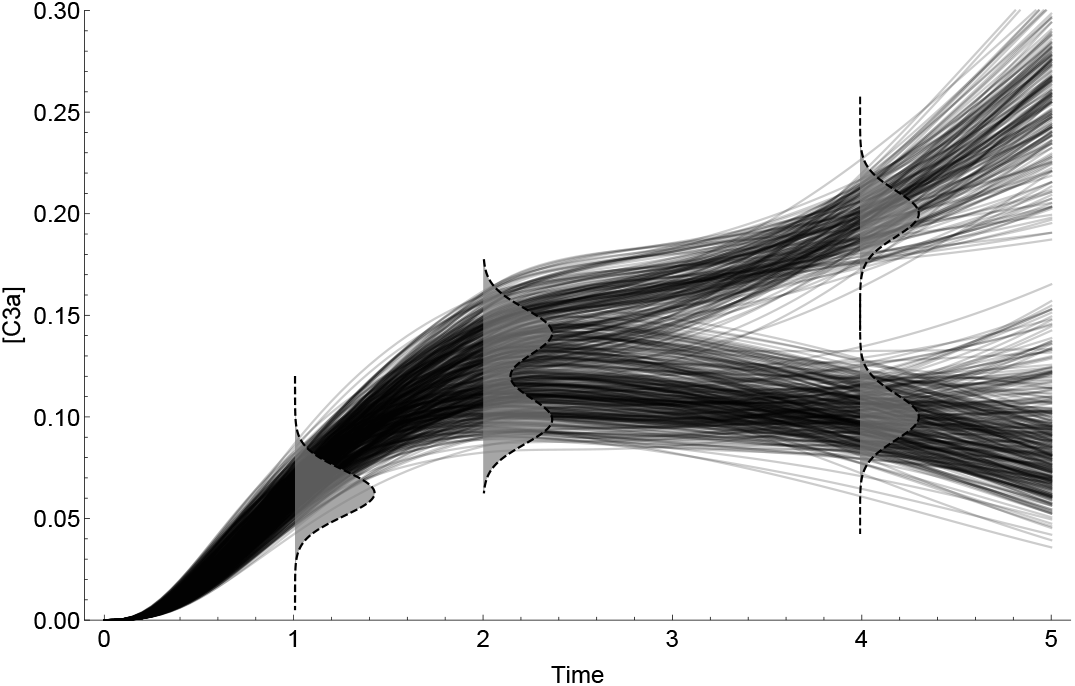
TNF signalling pathway model. **Target output distribution (dashed plots with grey filling) and unique trajectories (black solid lines) obtained from the posterior parameter distribution**. In CMC, 10,000 independent samples were used in the “ContourVolumeEstimator” step, and 5,000 MCMC samples across each of 4 Markov chains were used in the second, with the first half of the chains discarded as “warm-up” [21].

## 5 Discussion

Determining the cause of variability in cellular processes is crucial in many applications, ranging from bioengineering to drug development. In this paper, we introduce a Bayesian method for estimating cellular heterogeneity from “snapshot” measurements of cellular properties, taken at discrete intervals during experiments. Our approach assumes what we call a “heterogeneous ordinary differential equation” (HODE) framework, in which biochemical processes in all cells are governed by a common ODE. In HODEs, each cell has different rate parameter values, causing a variety of measurements to be obtained across cells. In this framework, estimating heterogeneity in cellular processes amounts to determining the probability distributions of parameter values of the governing ODE. Our method of estimation is a two-step Monte Carlo sampling process we term “Contour Monte Carlo” (CMC), which does not require the number of cell clusters to be provided before estimation, unlike for other approaches. CMC can be used to process high volumes of individual cellular measurements since the framework involves fitting a kernel density estimator to raw experimental data and using these distributions rather than data as the target outcome. CMC can handle arbitrary multivariate structure in measured outputs, meaning it can capture correlations between the same cellular species at different timepoints or, for example, contemporaneous correlations between different cellular compartments. Being a Bayesian approach, CMC uses prior distributions over parameter values to ensure uniqueness of the posterior distribution, allowing pre-experimental knowledge to be used to improve estimation robustness. The flexible and robust framework that CMC provides means it can be used to perform automatic inference for wide-ranging systems of practical interest.

Our approach also provides a natural way to test that the process is working satisfactorily. Feeding posterior parameter samples obtained by CMC into forward model simulations results in a distribution of output values which can be compared to the target. Indeed, we have found this comparison indispensable in applying CMC in practice and include it as the last step in the CMC algorithm (Algorithm 1). Discrepancies between the target output distribution and its CMC approximation can occur either as a result of poor estimates of the “contour volume distribution” in the first stage of the algorithm or due to insufficient MCMC samples in the second. Either of these issues are often easily addressed by increasing sample sizes or changing hyperparameter settings for the kernel density estimator. Although kernel density estimation in high dimensional spaces remains an open research problem, we have found vine copula kernel density estimation works well for the dimensionality of output measurements we investigate here [23].

Failure to reproduce a given output distribution can also indicate that the generating model (the priors and the forward model) are incongruent with experimental results. This may either be due to misspecification of the ODE system or because the assumption of a deterministic forward model is inappropriate. Our approach currently assumes that output variation is dominated by cellular variation in the parameter values of the underlying ODE, with measurement noise making a negligible contribution. Whether this is a reasonable assumption depends on the system under investigation and, more importantly, on experimental details. We recognise that neglecting measurement noise when it is, in fact, important in determining observed data means CMC will overstate cellular variation. It may also mean that some output distributions cannot be obtained by our model system (i.e. HODEs without noise). Future work incorporating a stochastic noise process or, more generally, including stochastic cellular mechanisms is thus likely to be worthwhile.

We have labelled our approach as Bayesian since it involves explicit estimation of probability distributions and requires priors. We recognise, however, that it is not of the form used in traditional Bayesian inference. This is because, rather than aiming to formulate a model that describes output observations, our approach aims to recapitulate output *distributions*. Others [20], (including us [22]), have considered similar problems before; perhaps most notably by Albert Tarantola in his landmark work on inverse problem theory (see, for example, [35]). In Tarantola’s framework, a joint input parameter and output space is considered, where prior knowledge and experimental theory combine elegantly to produce a posterior distribution whose marginal output distribution is a weighted “conjunction” of various sources of information. This work has seen considerable interest in areas such as the geosciences [36, 37], and we propose that Tarantola’s approach may prove useful for the biosciences.

The natural world is rife with variation, and mathematical models represent frameworks for understanding its causes. Typically, the state of biological knowledge is such that one effect - a given pattern of variation - has many possible causes. Observational or experimental data can be used to apportion weight to each cause, in a process that amounts to solving an inverse problem. The approach we describe here follows the Bayesian paradigm of inverse problem solving where uncertainty in potential causes (i.e. parameter values) is described using probability distributions. Here, we illustrate the worth of our method by using it to estimate cellular heterogeneity in biochemical processes. However, it could equally be used to invert other classes of under-determined systems arising elsewhere. Contour Monte Carlo provides an automatic framework for performing inference on such under-determined systems, and the use of priors allows for robust and precise parameter estimation unattainable through the data alone.

## 6 Author contributions

BL, DJG and SJT conceived the study. BL carried out the analysis. All authors helped to write and edit the manuscript.

